# Edge-enhanced disruptive camouflage is contrast dependent in VR environments

**DOI:** 10.64898/2026.10.01.755649

**Authors:** Ioan E. Smart, Rebecca J. Sharman, Kenneth C. Scott-Brown, P. George Lovell

## Abstract

In camouflage research, field studies provide ecological validity but lack experimental control, and computer-based experimental studies have the opposite issue. Virtual reality (VR) systems could potentially allow researchers to have both experimental control and ecological validity. Here we develop a VR environment using Unreal Engine and present it via a head-mounted display (HMD). The HMD affords binocular depth and therefore a more ecologically valid, three-dimensional environment, whilst still maintaining experimental control. Disruptive camouflage interferes with depth-based cues, making it a good candidate for testing within a VR environment. Disruptively camouflaged moth-like targets were presented to human surrogate predators, and their survivability was measured. We show that VR can be used to measure camouflage effectiveness by replicating existing findings showing that edge-enhanced disruptive camouflage (EE-DC) is more effective than disruptive camouflage (DC) alone. Furthermore, we extend these findings to show that the effectiveness of EE-DC is modulated by the relative contrast of the disruptive patches. Specifically, it is more effective when the contrast difference between the patches is low. Additionally, we considered edge-enhancement in isolation and found that it still provided a camouflage benefit even without a DC component. Search paradigms within an immersive VR experimental framework offer a unique opportunity to evaluate camouflage theory in a rigorous, yet ecologically grounded manner.

**Significance Statement:** When trying to measure camouflage effectiveness, researchers currently have two main methods: field studies, and computer-controlled experiments. However, both have limitations; field studies are very realistic, but difficult to control, whereas laboratory studies are highly controlled, but not very realistic. We propose that using a virtual reality (VR) paradigm would give researchers the best of both worlds. Using VR, we replicated existing findings and collected new data showing the importance of contrast in specific types of camouflage effectiveness. This establishes a methodological advance: researchers can now act as “digital architects” and control the visual information present in the environment. This flexibility makes it possible to investigate both longstanding and emerging questions about visual perception within ecologically realistic, experimentally controlled environments.

## Introduction

There are currently two key methods to measure behavioral responses to camouflage patterns: 1) laboratory-based studies, using mainly psychophysics-style computer-based experiments with humans (1–5); 2) field-based studies based upon survival-analysis, where predation levels of animals are measured (3, 6, 7). One can consider the two methods to lie on a continuum, with computer-based psychophysical methods having a high level of experimental control, but low ecological validity. Whereas field-based studies have higher ecological-validity at the expense of control. To date, there is no one technique that can truly offer high validity and control. We propose that virtual reality (VR) may offer a solution to this problem.

Efforts have been made to develop methodologies that allow for both high control and high validity; however, current methods still retain significant limitations. Wheatstone stereoscopes allow for binocular-stereopsis in computer-based experiments, but the field of view is limited, and participants cannot make naturalistic head-movements (8). Careful weather monitoring can increase control in field studies but it cannot prevent natural variation over time (9). Significant strides have been taken to incorporate real-world complexity into both types of experimental design. For example, behavioral effects like distance dependence (10, 11) and orientation of targets in relation to background textures (12–16), and environmental factors like dynamic illumination (17–19). However, these studies are all still subject to an inherent trade-off between experimental control and ecological validity. Bian and colleagues (20–22) used 3D-animated virtual environments to model animal behavior under a range of environmental conditions. However, these environments were only used for computational modelling rather than any behavioral measures. These studies highlight the value of cross-discipline research, principally incorporating biological and computational science (see 23). VR systems have already been used successfully to examine visual search processes demonstrating their potential (24–26). However, animal camouflage research has not yet been able to acquire behavioral data in a controlled immersive 3D environment (see 27 Section 4, for a review on camouflage in 3D).

Contemporary VR systems simulate physical presence through a head mounted display (HMD) with a wide field of view (FoV), which allows real-time interactions through motion tracking (see 28 for a review). By presenting separate images to each eye, VR HMDs introduce binocular disparity, enabling stereoscopic depth perception. This is particularly important to camouflage research because stereoscopic depth has been postulated as an important camouflage breaking mechanism (29), but a majority of methodologies have not allowed testing of this hypothesis. The few studies that have tackled this question have been limited by small screen size (as mentioned above 8). VR systems, however, provide immersive environments with the potential for 360° search areas. This is important because animal camouflage is designed to operate in a wider environment that spans the full visual field. Experiments that utilize relatively small search areas narrow the potential locations of the target to regions within the fovea or near-periphery regions, where acuity is relatively high. In contrast to lab-based studies, targets in the real-world may spend much of their time only visible to relatively poor peripheral vision; while optimal coloration and patterning for concealment may be different for the periphery and fovea. Therefore, it is difficult to make inferences from traditional psychophysical experiments to the real-world. It is possible that the nature of a screen-based search task might be very different to the equivalent real-world scenario, due to the fundamentally different perceptual demands. VR provides the opportunity to design experiments involving freely moving observers interacting with a 3-dimensional world (30). It offers naturalistic features such as occlusion and parallax (e.g., 31), weather, and visual clutter, along with a field of view almost as large as that available in the real world.

Disruptive camouflage impedes both detection (1, 32–34) and recognition (4, 5) by using high-contrast patches (maximal disruptive contrast) positioned at the body’s edge to break up its outline (marginal markings) or on particularly recognizable features (coincident disruption). These patches create false edges and visual noise (surface disruption), which segment the organism into components that appear to belong to different depth planes (differential blending) (35). These five features, maximal disruptive contrast, marginal marking, coincident disruption, surface disruption, and differential blending, comprise the principles of disruptive coloration and underpin our understanding of its effectiveness.

In nature, the false edges generated from disruptive camouflage patterning are sometimes accentuated through a phenomenon known as edge-enhancement; dark patches have darker edges and light patches have lighter edges (1, 4, 5, 36). This is characterized by a sawtooth pattern of luminance at the intersection of the patches. It is proposed that edge-enhancement facilitates camouflage by increasing the saliency of the false edge, detracting the perceiver away from the true outline, and by emphasizing false depth cues (35). Disruptive camouflage is of particular interest for VR methods because it is proposed to impede detection by interfering with depth-based cues (37). Recent studies have suggested that the benefits afforded by edge-enhancement are eliminated by binocular-stereopsis (8). Specifically, the researchers suggested that reliable depth information provided by binocular disparity over-ruled the depth boundaries implied by edge-enhancement. Here we will investigate whether the depth cues created in VR will undermine any crypsis afforded by disruptive coloration and edge-enhancement.

In the current study, we show that VR provides a powerful means of conducting ecologically valid, highly controlled research in camouflage, revealing interactions between subtle pattern characteristics and visual perception. We examine the effectiveness of edge-enhanced disruptive camouflage and investigate whether the principles of differential blending, surface disruption, and maximal disruptive camouflage (37, 38) hold in a virtual reality environment. In addition, we explore whether edge enhancement can increase disruptive camouflage effectiveness, and whether it influences crypsis in isolation. The design took inspiration from the survival type experiments where predation is examined across an interval of time. This design was originally created by Cuthill et al (32), and has subsequently been implemented in other camouflage research (2, 3, 6, 7, 39, 40). We use humans as surrogate predators and examine their ‘predation’, *i.e., ‘*shooting’ of a single target, within a 30 second trial window for nine types of camouflaged targets. The camouflaged patterns on the targets (Pattern) were comprised of nine conditions: a no/low/high contrast Disruptive camouflage (DC) component and no/low/high contrast Edge-enhancement (EE) component, e.g. DCHiEELo see Figure 1. We predict that:

1. In accordance with the principles of differential blending and surface disruption, targets with disruptive patterning will have higher survival than a uniform pattern.
2. In accordance with the principle of maximal disruptive contrast, a large contrast difference between the disruptive patches will increase the survivability of a disruptively patterned target.
3. Edge-enhancement will increase camouflage effectiveness compared to DC alone as the extended search space will mean that the targets are not always examined binocularly, and subsequently the benefits afforded by EE should remain.
4. Edge-enhancement in isolation will increase camouflage effectiveness compared to a uniform pattern by introducing false edges that interfere with the organism’s true outline.

**Figure 1.**
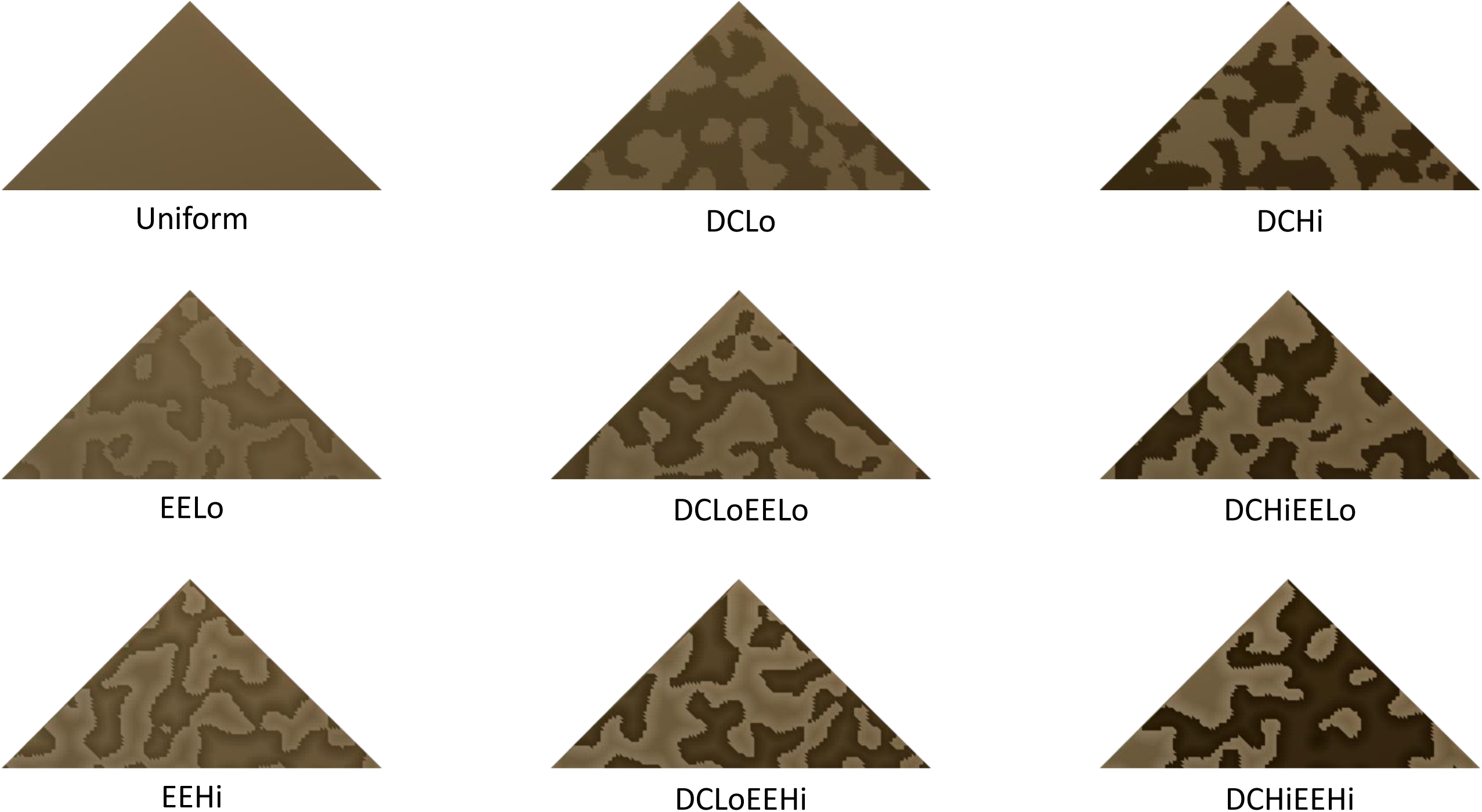
The 9 pattern types. Conditions that were ‘Lo’ had a contrast-step of 1, which was approximately 7.88 L*, whereas the ‘Hi’ conditions had a contrast-step of 2. Note that for the EE conditions the offset corresponds with ± ½ a contrast step. See Figure 4 for examples of these patterns in-situ.

## Results

Data and analysis files are available through OSF (41).

### Statistical analyses

Statistical analyses were performed using R studio (R foundation for Statistical Computing, www.R-project.org version 2024.12.1). Survival data were processed using the “survival”, “survminer”, and “coxme” packages (42–44), and analyses proceeded in three stages. First, proportional hazards (PH) models were used to identify an appropriate fixed-effects structure for subsequent modelling. Second, because Pattern represents the full 3×3 combination of DC and EE contrasts, a separate PH model including Pattern and stimuli presentation order was used to obtain descriptive pattern-level hazard ratios and Kaplan-Meier curves. Finally, mixed-effects Cox models incorporating a random intercept for participant were fitted to provide the inferential tests of DC, EE, and their interaction.

A total of 2672 out of 3600 targets (74.22%) were detected and ‘predated upon’ within the 30 second trial window. The rest were treated as censored in the survival analysis.

PH models were fitted without random intercepts because diagnostics for evaluating PH assumptions are not available for mixed-effects Cox models. These PH models were used to determine the fixed effects structure and justify the following component-wise mixed effects analysis. These models included the DC contrast component, the EE contrast component, their interaction, and the stimulus presentation order as fixed effects:

1. ∼ DC_contrast*EE_contrast + Order, cluster =Participant
2. ∼ Order, cluster =Participant
3. ∼ DC_contrast*EE_contrast, cluster =Participant
4. ∼1, cluster =Participant

Model 1 fit the data best (AIC = 40116.02), with all simpler models showing poorer fit (ΔAIC > 144). Schoenfeld residuals indicated a mild proportional-hazards violation for the DC contrast component, reflecting a time-varying effect of DC within trials (i.e., DC has a stronger influence early in the search window). No major departures from proportionality were detected for EE contrast or Order. As these PH models were used only to identify the fixed-effects structure and to justify the subsequent pattern-level descriptive analysis, the mild DC violation did not alter the selected structure.

As Pattern represents the full 3×3 combination of DC and EE contrast, a separate PH model including Pattern and Order was used to obtain descriptive pattern-level hazard ratios and to create plots.

Within the 30 second trial window more than fifty percent of the Uniform pattern are ‘predated upon’ within the first 5 seconds. Fifty percent of EELo, EEHi, DCHiEEHi, DCLo are predated within 10 seconds. It took at least 15 seconds for 50% of DCHiEELo and DCHi and 25 seconds for DCLoEELo to be predated upon. Finally, less than 50% of the DCLoEEHi pattern were predated upon within the 30 second trial window. These results are displayed in Supplementary Table 1 and a breakdown of the predation events, at every 5 second interval, for each pattern, are presented in Figure 2. Patterning with disruptive elements show approximately 58 – 81% higher survival than Uniform patterning (see Figure 3 and Supplementary Table 2). Additionally, EEHi shows a 40% higher survival than Uniform patterning (HR = 0.60, 95% CI = 0.52 - 0.69).

**Figure 2.**
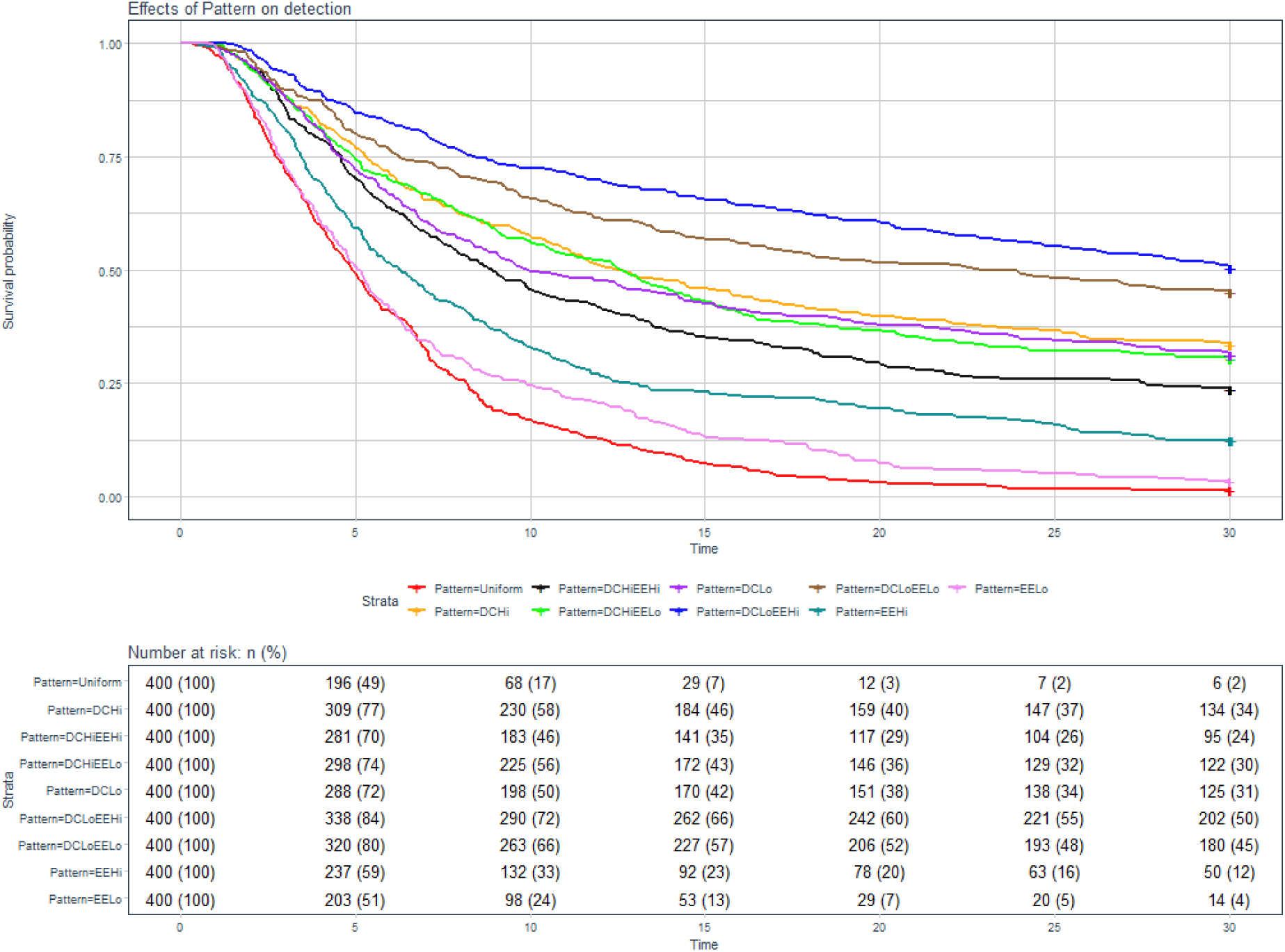
Survival curves of pattern treatments. Curves indicate the probability of avoiding ‘predation’ as a function of time over the 30 second predation window, based on Kaplan-Meier estimates to account for incomplete data owing to censoring. The table indicates the number (and absolute percentage) of targets at risk at every 5 second interval of the ‘predation window’.

**Figure 3.**
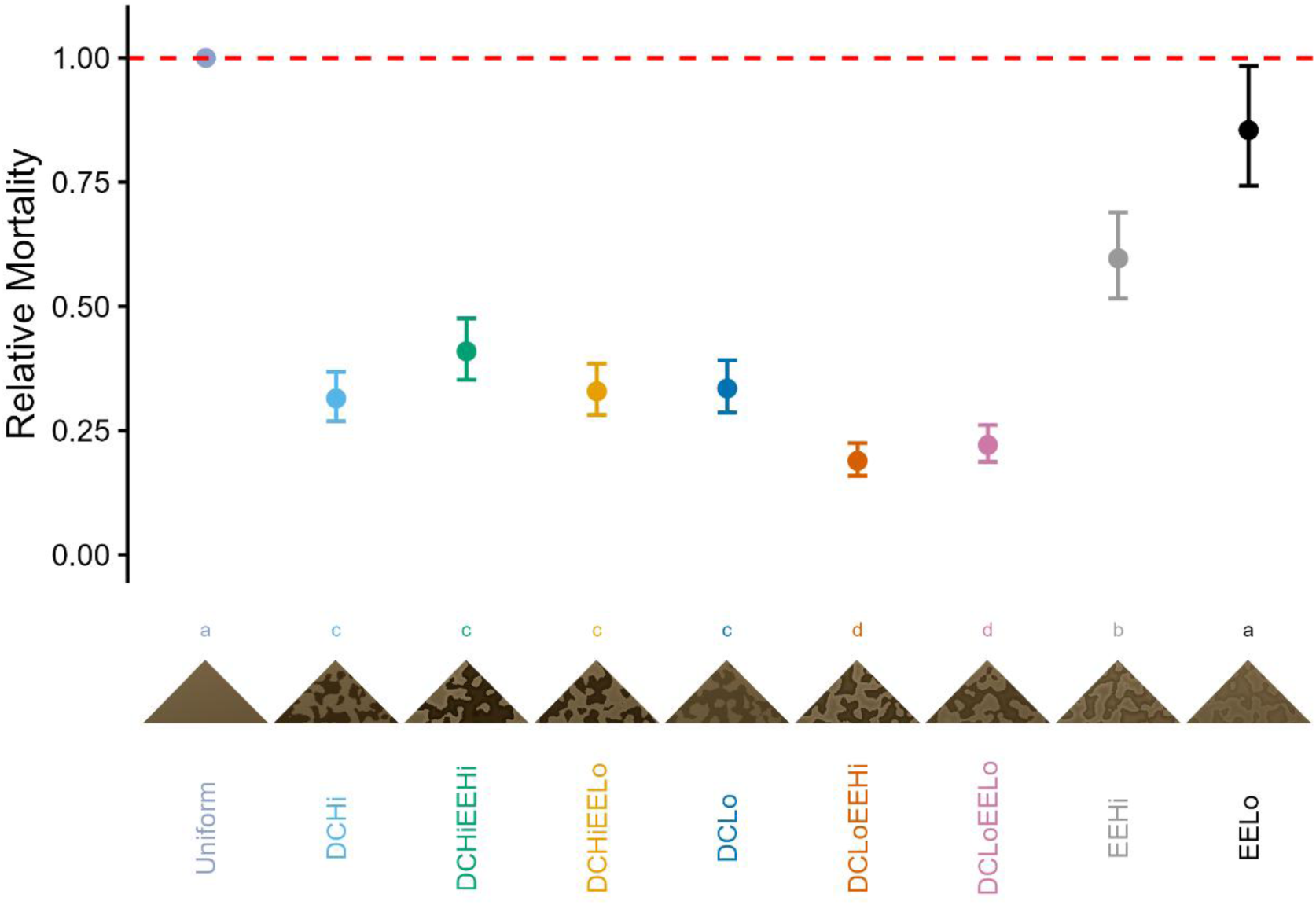
The hazard ratios, with 95% confidence intervals, for each pattern condition (N = 20) with reference to Uniform patterning. Groups sharing the same superscript (a - d) have no significant difference from each other in the post hoc Bonferroni adjusted comparisons.

Bonferroni adjusted, Tukey-type, pairwise comparisons (R package multcomp 45) indicated Uniform and EELo were significantly easier to detect and shoot than all others (see Table 1). EEHi had intermediate survival, but significantly worse survival than DCHi (z = 7.84, p < 0.001). Furthermore, DCHi was not significantly different from DCLo (z = 0.72, p = 1.000), DCHiEELo (z = 0.52, p = 1.000), DCHiEEHi (z = 3.14, p = 0.061). Moreover, DCHi had significantly worse survival than DCLoEELo (z = 3.88, p = 0.004) and DCLoEEHi (z = 5.43, p < 0.001).

**Table 1.** Bonferroni adjusted Tukey-type pairwise comparisons from the proportional hazards model fit with pattern. Values toward the right of the diagonal correspond with Bonferroni-adjusted p-values, whilst values toward the left correspond with Tukey z-statistics. Negative z-values indicate that the row survived better than the column treatment.

|  | *Uniform* | *EELo* | *EEHi* | *DCLo* | *DCLoEELo* | *DCLoEEHi* | *DCHi* | *DCHiEELo* | *DCHiEEHi* |
| --- | --- | --- | --- | --- | --- | --- | --- | --- | --- |
| *Uniform* |  | 1.000 | <0.001 | <0.001 | <0.001 | <0.001 | <0.001 | <0.001 | <0.001 |
| *EELo* | -2.192 |  | <0.001 | <0.001 | <0.001 | <0.001 | <0.001 | <0.001 | <0.001 |
| *EEHi* | -7.000 | -4.865 |  | <0.001 | <0.001 | <0.001 | <0.001 | <0.001 | <0.001 |
| *DCLo* | -13.740 | -11.784 | -7.150 |  | <0.001 | <0.001 | 1.000 | 1.000 | 0.556 |
| *DCLoEELo* | -17.647 | -15.847 | -11.495 | -4.594 |  | 1.000 | 0.004 | <0.001 | <0.001 |
| *DCLoEEHi* | -18.813 | -17.076 | -12.860 | -6.125 | -1.589 |  | <0.001 | <0.001 | <0.001 |
| *DCHi* | -14.384 | -12.445 | -7.842 | -0.720 | 3.879 | 5.425 |  | 1.000 | 0.061 |
| *DCHiEELo* | -14.005 | -12.042 | -7.385 | -0.201 | 4.415 | 5.954 | 0.522 |  | 0.303 |
| *DCHiEEHi* | -11.593 | -9.556 | -4.796 | 2.422 | 6.967 | 8.453 | 3.138 | 2.635 |  |

As an overview Tukey-type pairwise indicate: **Uniform = EELo < EEHi < DCHiEEHi = DCLo = DCHiEELo *=* DCHi < DCLoEELo = DCLoEEHi;** see Table 1 and Figure 3.

Odds Ratios (R package epitools 46) support this, showing that EELo has an equivalent level of survival to Uniform pattern by the end of the trial (OR = 2.34, 95% CI = 0.92 – 6.79, χ^2^ *=* 0.070). All other patterns were significantly more likely to survive than Uniform patterning (Supplementary Table 3; Supplementary Figure 1).

Exploratory plots indicated the effect of stimuli presentation order across the course of the experiment. Data indicates that participants predate upon more targets as the experiment progresses (Supplementary Figure 2). Plotting response time against stimuli presentation order for the successful predation data shows that participants are quicker to respond as the experiment progresses (Supplementary Figure 3). Finally, plotting response time against stimuli presentation order for *all* data shows that participant response times decrease as the experiment progresses (Supplementary Figure 4). These plots suggest that participants were getting better at detecting targets.

To identify which DC and EE contrast components drive the observed pattern differences, we fitted mixed-effects Cox models with DC contrast, EE contrast, and their interaction as fixed effects. These models included a random intercept for participants to account for repeated measures and used the fixed-effects structure previously supported by the PH models.

1. ∼ DC_contrast*EE_contrast + Order + (1|Participant)
2. ∼ Order + (1|Participant)
3. ∼ DC_contrast*EE_contrast + (1|Participant)
4. ∼1 + (1|Participant)

Model 1 fits the data best (AIC = 39495.06), with all simpler models showing poorer fit (ΔAIC > 16). Both DC contrast and EE contrast showed significant main effects on hazard (DC_Cont1: HR = 0.30, z = –14.97, p < 0.001; DC_Cont2: HR = 0.28, z = –15.79, p < 0.001; EE_Cont1: HR = 0.86, z = –2.18, p = 0.029; EE_Cont2: HR = 0.57, z = –7.66, p < 0.001; Table 2). Several DC*EE interactions were also significant (see Table 2). DC_Cont1:EE_Cont1 showed a synergistic reduction in hazard (HR = 0.73, z = –2.76, p = 0.006), whereas DC_Cont2:EE_Cont2 showed strong interference (HR = 2.37, z = 7.67, p < 0.001). The remaining interactions were either weak (DC_Cont2:EE_Cont1: HR = 1.24, z = 1.91, p = 0.056) or non-significant (DC_Cont1:EE_Cont2: HR = 0.92, z = –0.67, p = 0.50), indicating that the hazard for these combinations was well explained by the additive main effects of DC and EE.

**Table 2.** Output from the mixed-effects Cox model, decomposing patterns into DC and EE contrast components. Note that exp_coef correspond with hazard ratios (HRs). A reduced hazard (improved survivability) is indicated by a HR < 1, whereas an increased hazard (poorer survivability) is indicated by a HR > 1. Interaction terms quantify whether specific DCxEE combinations show synergy (HR < 1) or interference (HR > 1) relative to the additive main-effects model.

|  | *coef* | *exp_coef* | *se_coef* | *z* | *p* |
| --- | --- | --- | --- | --- | --- |
| *DC_Cont1* | -1.201 | 0.301 | 0.080 | -14.970 | < 0.001 |
| *DC_Cont2* | -1.279 | 0.278 | 0.081 | -15.790 | < 0.001 |
| *EE_Cont1* | -0.157 | 0.855 | 0.072 | -2.180 | 0.029 |
| *EE_Cont2* | -0.568 | 0.567 | 0.074 | -7.660 | < 0.001 |
| *Order* | 0.005 | 1.005 | 0.000 | 12.660 | < 0.001 |
| *DC_Cont1:EE_Cont1* | -0.319 | 0.727 | 0.116 | -2.760 | 0.006 |
| *DC_Cont2:EE_Cont1* | 0.214 | 1.239 | 0.112 | 1.910 | 0.056 |
| *DC_Cont1:EE_Cont2* | -0.079 | 0.924 | 0.119 | -0.670 | 0.505 |
| *DC_Cont2:EE_Cont2* | 0.861 | 2.366 | 0.112 | 7.670 | < 0.001 |

These component-wise effects clarify the pattern hierarchy observed in the descriptive analysis: patterns with low DC and low EE (i.e., DCLoEELo) benefit from synergy, patterns with low DC and high EE (i.e., DCLoEEHi) show strong additive benefits, and patterns combining high DC with high EE (i.e., DCHiEEHi) show interference, reducing survivability relative to their main-effect predictions.

## Discussion

In accordance with the first prediction, disruptive patterning has higher survival than a uniform pattern. This aligns with existing results (6, 32, 47) and demonstrates that VR can be used to measure camouflage effectiveness. However, the second prediction was not met as contrast did not affect survival of disruptive coloration in the absence of edge-enhancement. The third prediction stated that edge-enhancement would provide substantial cryptic benefit, but this was only partially supported by the data. Adding edge-enhancement to disruptive coloration was only of significant benefit in the low contrast DC conditions, with the higher contrast DC equivalents failing to achieve the same effect. This could indicate that the advantages of edge-enhancement may be constrained by the relationship between contrast in the pattern and in the background. Exhibiting edge-enhancement independent of DC conditions did increase the survival over a uniform condition, and thus the fourth prediction was met.

It has long been postulated that stereopsis always acts as a camouflage breaking mechanism (29). Here we demonstrate that stereopsis does not necessarily eliminate the benefits of disruptive camouflage and edge enhancement. Despite participants having binocular cues available, pattern had a significant effect on detectability and survivability. Moreover, patterns with edge enhancement components significantly reduce detectability even further. This contrasts with findings that when binocular depth cues are available, stereopsis assists in the discrimination of false edges from real object boundaries, and consequently destroy any cryptic advantage provided by EE (8). However, this is in keeping with what we see in the natural world where many forms of (edge-enhanced) disruptive camouflage are widespread across taxa despite predation by binocular predators. Snakes, frogs and moths frequently exhibit (edge-enhanced) disruptive camouflage, despite their avian predators (e.g., *Agkistrodon contortrix contortrix*, *Limnodynastes tasmaniensis*, *Mimas tiliae*) (see Figure 1 in 1).

A major methodological difference between the current study and Adams and colleagues work (8) is the presentation method, specifically in terms of the field of view (FoV). In the current experiment the participant is situated in an environment that expands 360° around them, although the trees upon which the targets appear subtend only approximately 90° in the virtual space. In Adams and colleagues’ study (8) the full stimulus measured 39.64° by 28.74° visual angle. To contextualize this, the binocular field of view is ∼120° meaning our stimuli filled the majority of the visual field, and participants were required to move their heads and make eye movements to comfortably foveate all potential target locations and view them with high acuity. Stereopsis reduces with eccentricity (48); therefore, it is plausible that stereoscopic cues become less informative in a larger environment like ours, and instead participants need to serially/sequentially search the scene. The reduction in relative disparity, the larger FoV, and thus the need to search the scene may diminish the effects of stereopsis. Adams and colleagues (8) describe several instances in which the reliability of depth information drops. This includes large viewing distances, weak illumination, or when objects are partially occluded. The results indicate that under natural-looking immersive scenes displayed in VR, the influence of stereoscopic depth cues is reduced and the assumption that stereopsis breaks camouflage becomes questionable.

There are experimental differences, aside from presentation method, that may underpin the contradictory findings. Our targets existed on a depth plane similar to the background which reduces relative disparity depth cues needed for cyclopean stereopsis (48). However, the snake-like stimuli in Adams and colleagues study (8) also existed on depth planes similar to the background, including circumstances where the snake was not the closest object in the scene. It may be possible that the difference in the results is due to the shapes of the targets. In the current study, we replicated Cuthill and colleagues (6, 32) stimuli in VR and the moth target was completely flat whilst Adams and colleague’s study (8) involved a cylindrical snake-like target and thus exhibited curvature at the edges. A flatter target reduces the availability of curvature-based disparity cues, which may help preserve the effectiveness of EE.

In accordance with the principles of differential blending and maximal disruptive contrast, one would expect that patches that are maximally different from one another, whilst still existing within the range of the background colors, to be better camouflaged. This would increase the likelihood that the patches are not grouped together, but instead associated with the background, and disrupt perceptual organization (49). Therefore, as in our second prediction, one would expect high contrast DC to outperform low contrast DC, due to the increased contrast difference. However, our high and low contrast disruptive treatments were not significantly different in their detectability. Instead, both DC contrast levels had similar survival, indicating that within the luminance range used here, increasing DC contrast alone does not necessarily enhance camouflage (see Supplementary Figure 5 for the luminance range). It may be that a greater contrast difference is required to observe the effect, such as those used by Cuthill and colleagues (32).

In contrast to the DC conditions, there was a significant difference in detectability between the high and low contrast EE only conditions. Importantly, this shows that exhibiting EE on an otherwise uniform pattern will improve crypsis, as proposed by Egan and colleagues (1). However, the EE only conditions were not as successful at decreasing predation as the DC conditions. This may be due to the EE patterns being one uniform color, as opposed to two in DC, meaning that there is only one potential match for the background. The EE sawtooth luminance pattern does create a false edge, but it is unlikely that this would be as effective as the DC step-change. The sawtooth edge could fail to elicit Gestalt-like color grouping in the same way as DC coloration. This is particularly likely in the low-contrast EE only condition which did not significantly increase survival compared to the Uniform condition. The high-contrast EE only condition, whilst poorer than patterns with a DC element, had significantly better survival than the Uniform pattern. The greater contrast offset may have been sufficient to induce the Craik-Cornsweet illusion (50). This would cause the patches to appear different in their luminance, effectively creating an illusory DC pattern, resulting in the patches differentially blending with the background.

The introduction of EE to DC patterns is not simply additive, instead the contrast levels of the components interact, with some combinations producing synergy and others producing interference. When EE is added to a low contrast DC pattern the target becomes more camouflaged, but when it is added to a high contrast DC pattern it does not impact survivability. This suggests that the effectiveness of EE depends critically on the underlying DC contrast level. This is unlikely to be due to a difference in maximal disruptive contrast, and thus the saliency of the edge, as the contrast differences *within the targets* are equal (*e.g.,* DCHiEELo and DCLoEEHi, both have 3 contrast steps = 23.63L*). However, the underlying DC contrast affects the absolute value of the highest contrast points in the pattern. In the high-contrast DC patterns the addition of EE causes more extreme luminance values, some within the 0.5th percentile of luminance values in the background (Supplementary Figure 5). This would cause a failure in background matching, explaining the loss of crypsis in comparison to the lower-contrast DC patterns.

Many studies have investigated the principles of (disruptive) camouflage using static, non-immersive, environments. The presented paradigm merges the control attributed to computer-based detection experiments with the ecological validity of survival analysis approaches which are traditionally used to test camouflage theory. When searching the environment visually, the eyes and head work together (51) to scan the environment for valuable visual information that may indicate the presence of an object of interest (e.g., prey item). The head-mounted display and accompanying motion tracking, which enable real-time updates of a virtual environment to be displayed to the user, provide an opportunity to better scrutinize naturalistic visual search and camouflage. Importantly, this research offers a novel method for integrating ecological factors, such as habitat structure, sun positioning, environmental movement, and dynamic illumination, which influence camouflage effectiveness, within a controlled experimental framework. Our system lays the foundations for a new experimental paradigm that bridges controlled psychophysics and ecological realism, enabling rigorous, observer-dependent testing of camouflage principles under dynamic, real-world-inspired viewing conditions.

## Materials and Methods

### Participants

Opportunity sampling was used to recruit 20 students from Abertay University, two participants were authors (RJS and KSB). All participants had normal/corrected-to-normal vision. Ethical approval was obtained through the Ethics Committee at Abertay University (EMS4553). All participants were briefed and gave their informed written consent, in accordance with the Declaration of Helsinki.

### Apparatus

The experiment was conducted in a Meta Quest 2 device (52), with the associated right hand motion controller, and ran on a PC (Intel® Core™ i7-9700 CPU @ 3.00GHz; RAM 32.0GB; Nvidia GeForce RTX 2070 SUPER graphics card). The inter-pupillary distance (IPD) was set to 63mm, which is the average IPD of the adult population (53).

### Stimuli

Stimuli were created and presented using Unreal Engine 5 (version 5.0, 54). Within the application, the *X* axis represents the horizontal left/right plane, the *Z* axis represents the vertical up/down plane, and the *Y* axis represents the forward/backward plane. Objects within the application are scaled with unreal units (uu), where 1 uu corresponds with 1cm (55).

In each trial a single moth-target (a triangle: 25uu/2.24° wide, 12.5uu/1.12° tall; see Figure 4) would appear pseudo-randomly (*i.e.,* within a bounded space, see ‘Target location’ section) on one of 5 trees in front of the participant (see ‘The local background’ section). The moth-targets were comprised of two components: disruptive camouflage and edge enhancement.

**Figure 4.**
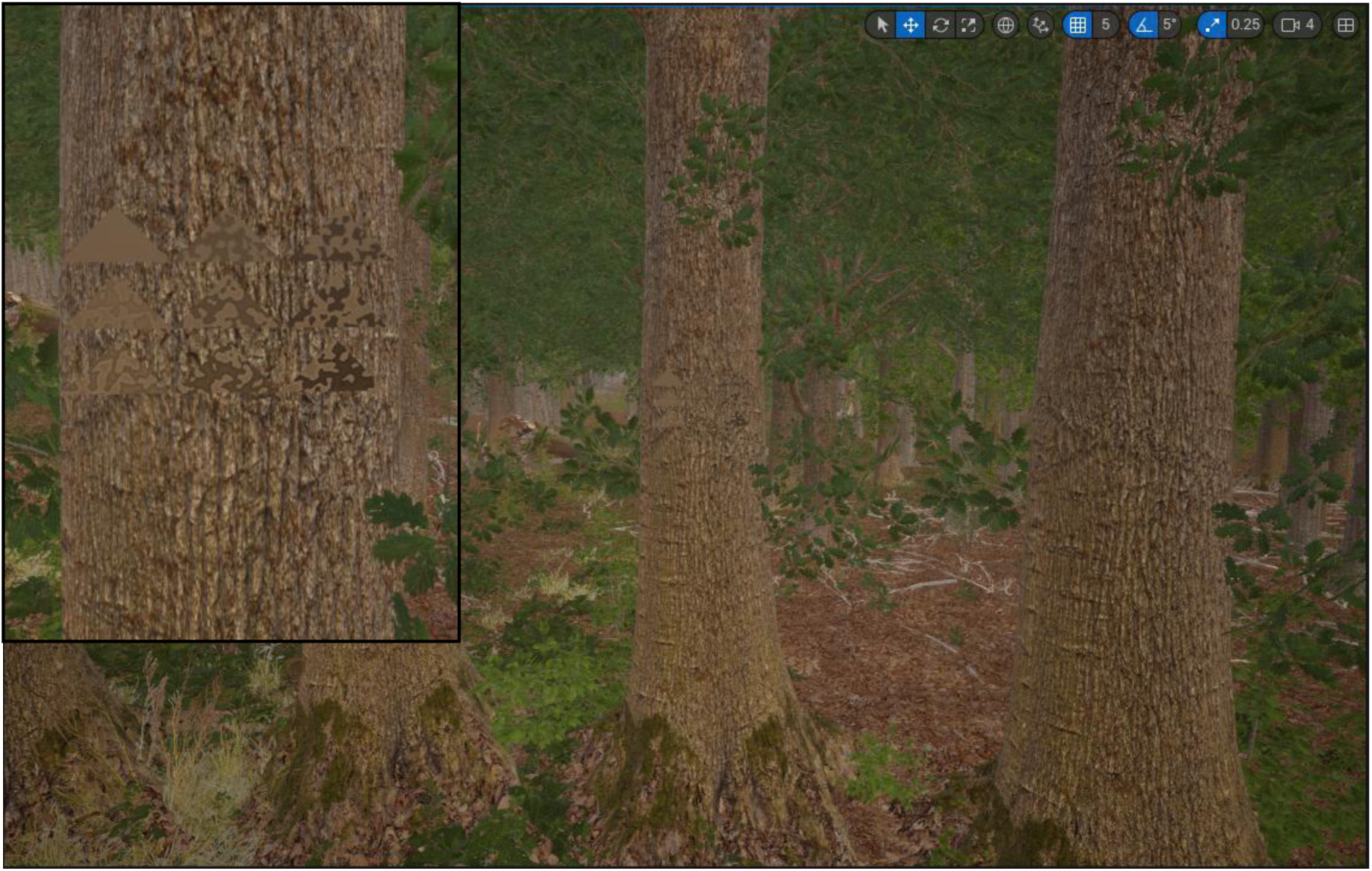
Examples of the 9 pattern types, see Figure 1, in-situ. The panel shows a zoomed in perspective of how the targets appear on the tree.

### Background Creation

#### The global background

Unreal Engine has a procedural foliage tool that enables the user to load in assets and select parameters for procedural generation (*e.g.,* max size or age of the foliage), such that a designated area can be populated by the parameterized assets (see 56).

Using assets downloaded from MAWI Global on the Unreal Marketplace (MW Broadleaf Forest, 57) a procedurally generated forest populated with European Beech (*Fagus sylvatica*) and German Oak *(Quercus pedunculata*) trees was created. A foliage blocking volume was placed around where the participant would be positioned so that foliage would not be generated too close to the participant. This also prevented the global background from occluding the local background upon which the targets would appear.

#### The local background

Whilst the procedurally generated forest created the global background, the immediate background upon which targets were positioned was controlled more precisely. Five trees, all exact replicas (Asset: SM_BF_BroadleafE_NoBase, 57), were placed equidistance from the participant at 750uu with a spacing of 22.5° (in virtual space) between them (Supplementary Figure 6). The rotation of the tree was randomized (within bounds) to minimize the unnatural look of duplicated trees. The randomization was bounded so that branches would not end up occluding moth targets.

### Lighting

The virtual environment was displayed under diffuse lighting and shadows were removed. Whilst this does make the environment more unnatural, or less-ecologically valid, it reduces confounding effects: (1) it prevents a different illumination gradient existing across the local backgrounds, which may result in location being a confounding factor for camouflage effectiveness; (2) shadows are great visual cues for 3-dimensionality (58). In pursuit of reducing variability in the data, these confounding factors were controlled in an effort to explicitly examine the effectiveness of the patterning. These confounds can be introduced in future experiments sequentially to examine the effect.

### Target creation

Using Matlab (R2019b; The MathWorks, Inc., Natick, MA, USA) we created a square trimesh and masked out a triangle, comprising 4950 smaller triangles, which would act as the artificial moth target. Camouflage for the targets was created in a similar way to that described in Egan *et al.* (1), where a white noise image is filtered in Fourier space to create a texture:

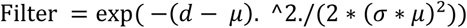

where *d* is the distance from the image centre (in Fourier-space), σ was 120 and µ was 0.06. Patches were assigned as a ‘HiColourPatch’ or the ‘LoColourPatch’ dependent upon whether they were in the upper or lower 50^th^ percentile of values in the filtered white noise image. The HiColourPatch corresponds with the modal colour present in the background, whilst the LoColourPatch corresponds with the contrast steps in the L* channel ***down*** from the HiColourPatch (1 contrast step = 7.88 L*, where 1 StdDev = 8.21). It was important to have one of the patches match the background, to adhere to the principle of differential blending (37). EE is determined in the most parsimonious way (sensu 1): offset and width. The contrast in DC is created by subtracting luminance from the modal colour; therefore, the EE offset is half the contrast step of DC to ensure that their contrast is equal. The width of EE corresponds with 0.09°. When the EE component is combined with the DC condition, this results in DCEE pattern conditions. If, however, the pattern only has an EE component and no DC, the HiColourPatch and LoColourPatch have equal values. After creation, these patterns were applied to the artificial moth target trimesh. Although moths are bilaterally symmetrical, this increases detection likelihood (2, 7, 39) and could confound disruptive effects, therefore, symmetrical patterning was not included.

There were nine pattern types that varied in their disruptive and edge enhancement components (see Figure 1 for an overview, Figure 4 for an example of these targets in-situ). To prevent participants becoming familiar with the texture of the camouflage pattern, we created each pattern twenty times. A pattern would appear on each of the five trees four times.

### Target location

On each trial, a target location was randomly chosen that falls upon the visible surface of the tree trunk. This was determined by taking a random coordinate from within a cuboidal plane and drawing a line trace toward a cylinder that encapsulated the tree (see Supplementary Figure 7). The point at which the line contacts the cylinder are the Cartesian coordinates at which the moth target would appear. To account for the curve of the tree, the rotation of the moth is set so that its surface is positioned at a tangent to the tree. Using a cylinder that is slightly wider than the tree and rotating the moth ensures that the whole surface of the moth target was visible and that there were no targets appearing inside the tree. Each target appeared on each tree four times, but the order the trees were selected was randomized. This meant that it was possible for targets to appear on the same tree successively.

### Task

Participants used the Quest 2 controller. The A button was used for interacting with menus (*e.g.,* starting the experiment, continuing the experiment after a pause, *etc.*). The grip button was used to show a laser, which is an aiming device. The trigger was used to ‘shoot’ the target. Participants were positioned within a procedurally generated forest environment and were tasked with detecting a moth-like target on one of five trees 750uu away. Upon detecting the target, the participant pressed and held a button to show a laser (to aim) and pulled the trigger to shoot at the target. This is the functional equivalent of moving a mouse to click on a target in a 2-dimensional (psychophysics-style) computer detection experiment. After correctly detecting and shooting a target, a confirmation sound played, the screen ‘blinked’ (*i.e.,* fade to black and back again in 1500ms), and a new trial started. If the participant was unable to locate the target within 30 seconds, the trial would move on automatically, and a sound played to indicate a ‘miss’. For each trial the location of the target on the tree, the rotation of the trees, and the patterning of the target were computed/loaded before the screen faded from black. Additionally, the position of the local background would jitter by ± 5.625° (in virtual space) trial to prevent rapid detection by pop-out, in situations when the targets appeared on the same tree as the previous trial. Prior to the experiment participants completed a set of practice trials (n = 27) to familiarize themselves with the ‘aim and shoot’ mechanisms of the experiment. The practice targets had a similar pattern to the test targets but were displayed at much higher luminance making the task much easier (*+ 10 L* on the test images*). After the practice and with the participants familiar with the controls/task they would start the main experiment, constituting 180 trials (maximum experiment duration = 75mins). During the experiment there were three instances where the experiment would pause, so that participants could remove the headset and take a brief break if desired.

## Author Contributions

I.E.S, P.G.L and R.J.S conceived and designed the experiment. I.E.S and P.G.L programmed the experiment; I.E.S. carried out the experiment and analysed the data; I.E.S. wrote the first draft of the manuscript with subsequent contributions from R.J.S, P.G.L. and K.S-B.

## Disclosure of Delegation to Generative AI

The authors declare the use of generative AI in the research and writing process. According to the GAIDeT taxonomy (2026), the following tasks were delegated to GAI tools under full human supervision:

– Code generation
– Code optimization

The GAI tool used was: Copilot.

Responsibility for the final manuscript lies entirely with the authors.

GAI tools are not listed as authors and do not bear responsibility for the final outcomes.

Declaration submitted by: Ioan E. Smart

Additional note: We used Copilot to process and optimize existing analysis of R code.

## Acknowledgments

Supported by an R-LINCS Studentship funded by Abertay University

## Supplementary Materials

**Supplementary Figure 1.**
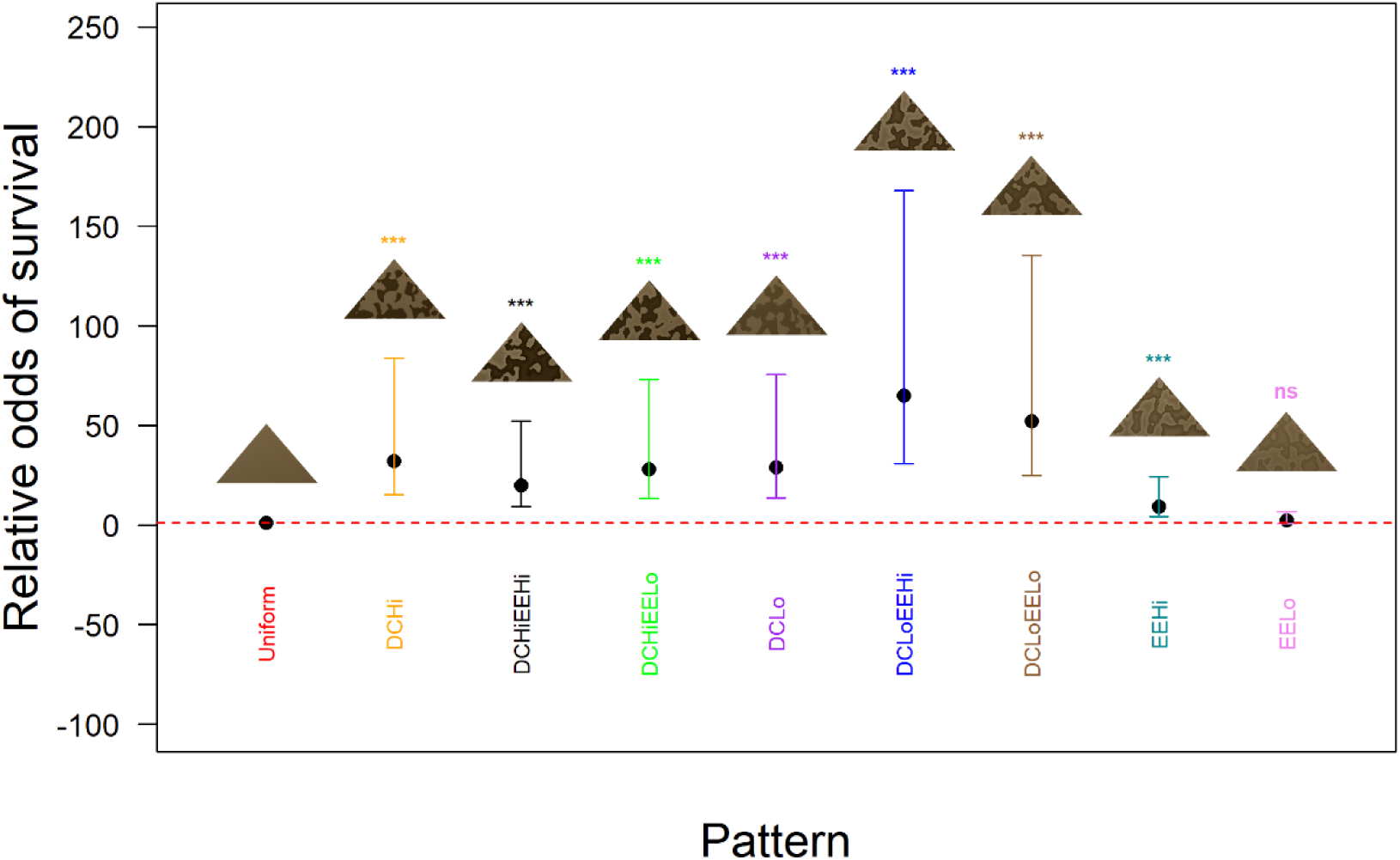
The odds ratios, with 95% confidence intervals, for each pattern condition (N = 20) with reference to Uniform patterning. Significance is indicated by *, where ‘***’ = p ≤ 0.001, ‘**’ = p ≤ 0.01, ‘*’ = p ≤ 0.05, and ‘ns’ = not significant.

**Supplementary Figure 2.**
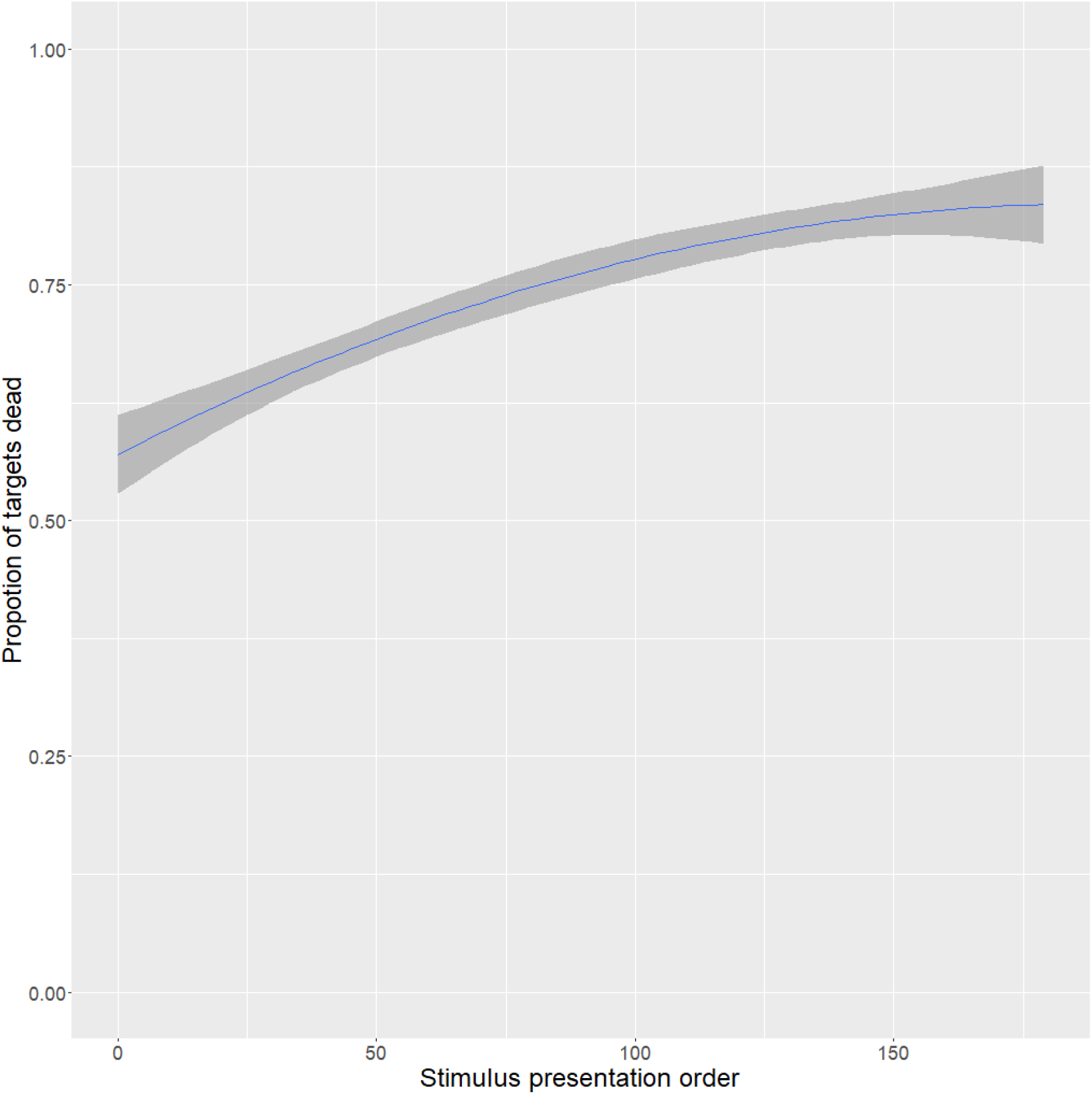
Proportion of targets successfully captured, with standard errors, as the experiment progresses.

**Supplementary Figure 3.**
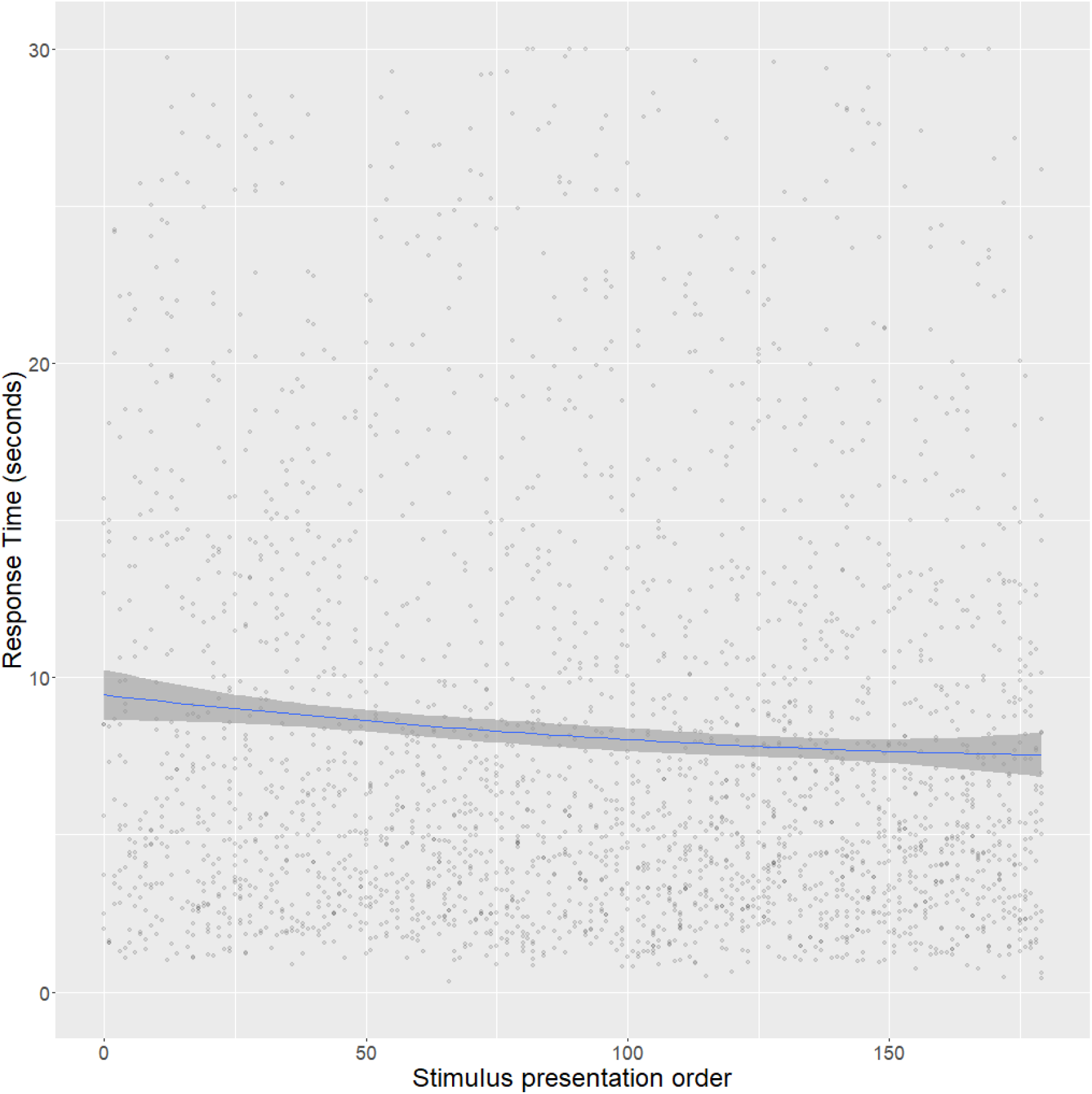
Response time, with standard errors, as the experiment progresses for captured targets.

**Supplementary Figure 4.**
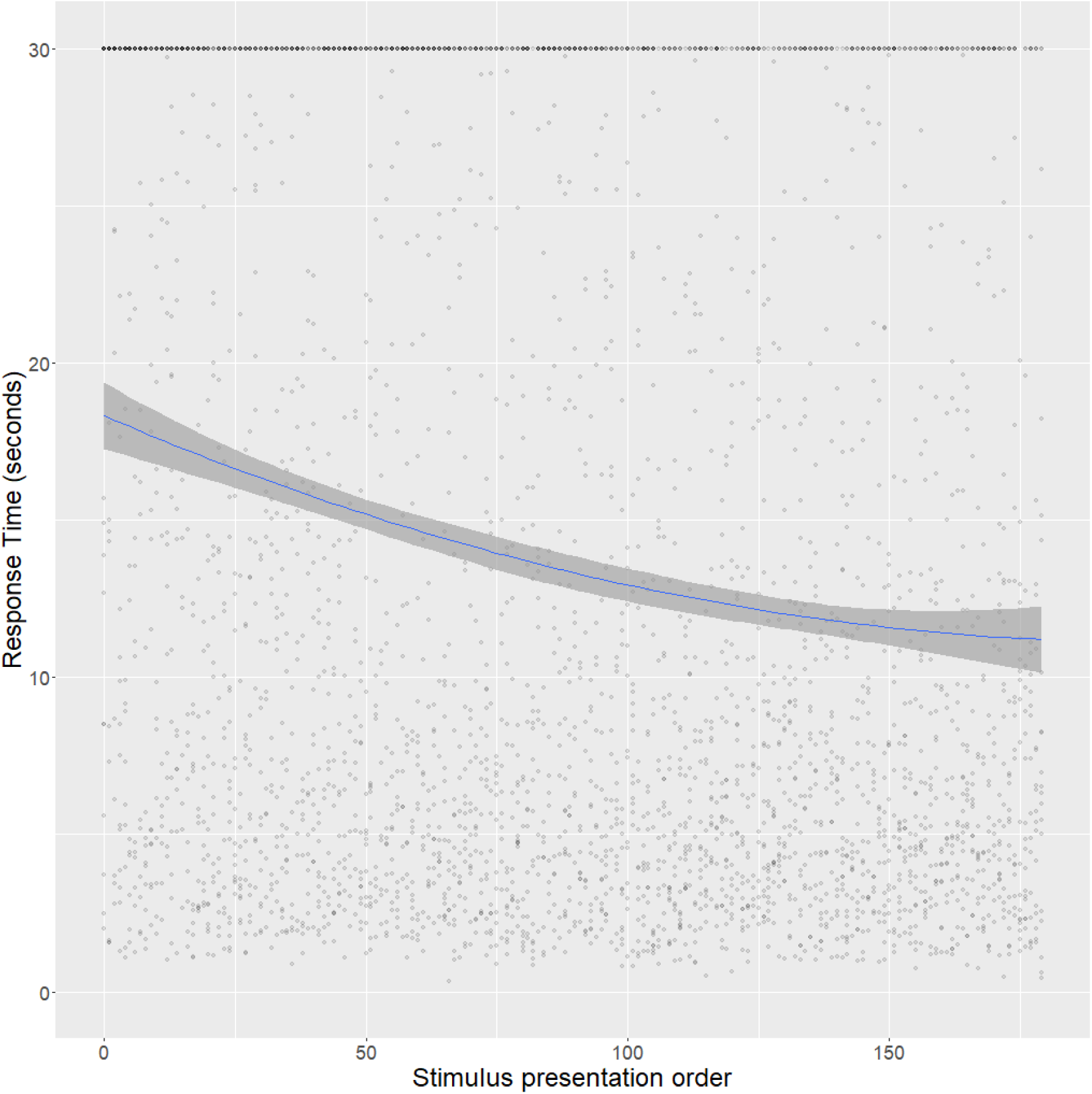
Response time, with standard errors, as the experiment progresses for all targets.

**Supplementary Figure 5.**
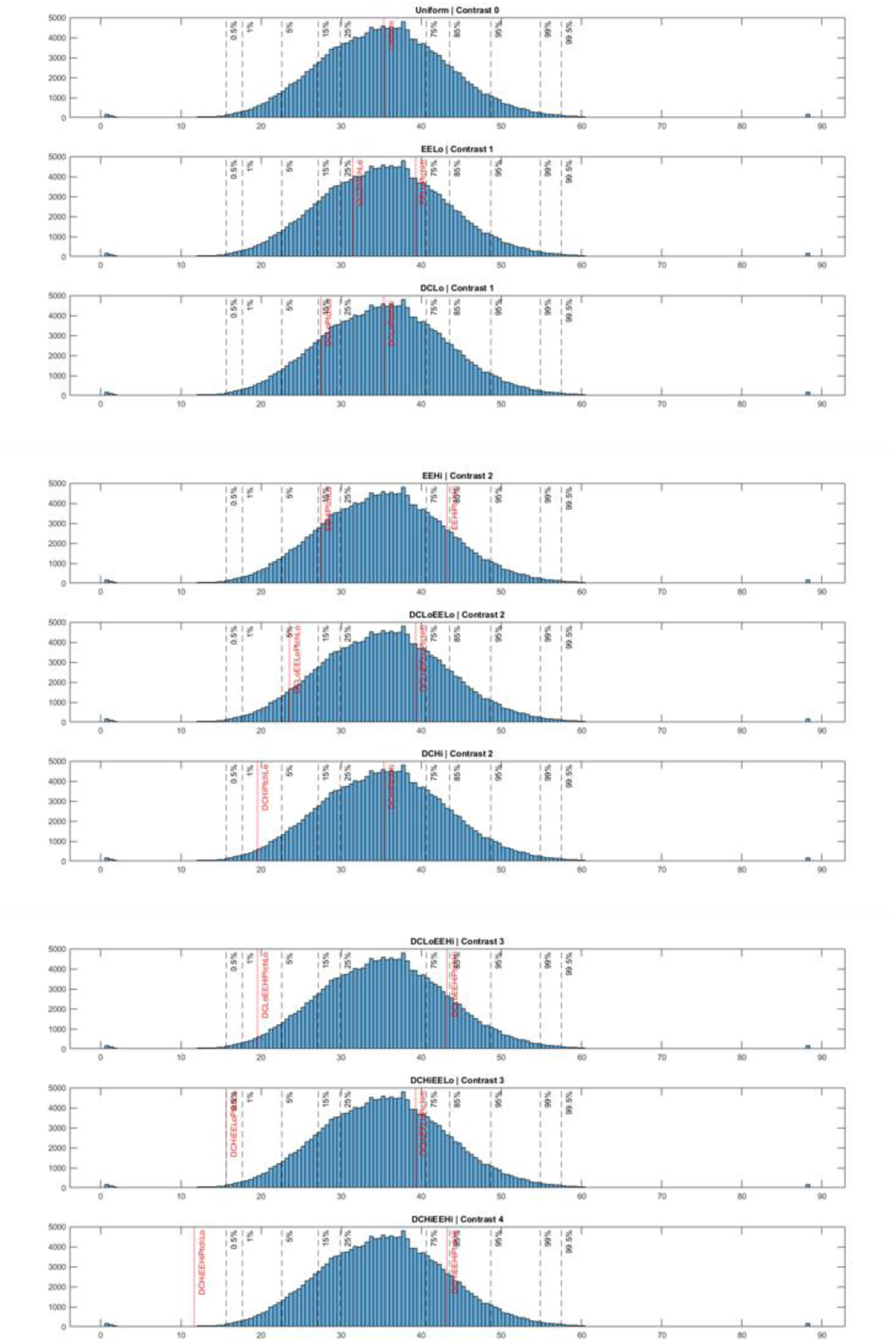
Luminance distribution (L*) for the background source image for stimuli creation overlaid with percentiles of L* values. Each panel corresponds with a pattern type with the red line indicating the max and minimum luminance. Dashed lines correspond with .5th, 1%, 5%, 15%, 25%, 75%, 85%, 95%, 99%, 99.5%.

**Supplementary Figure 6.**
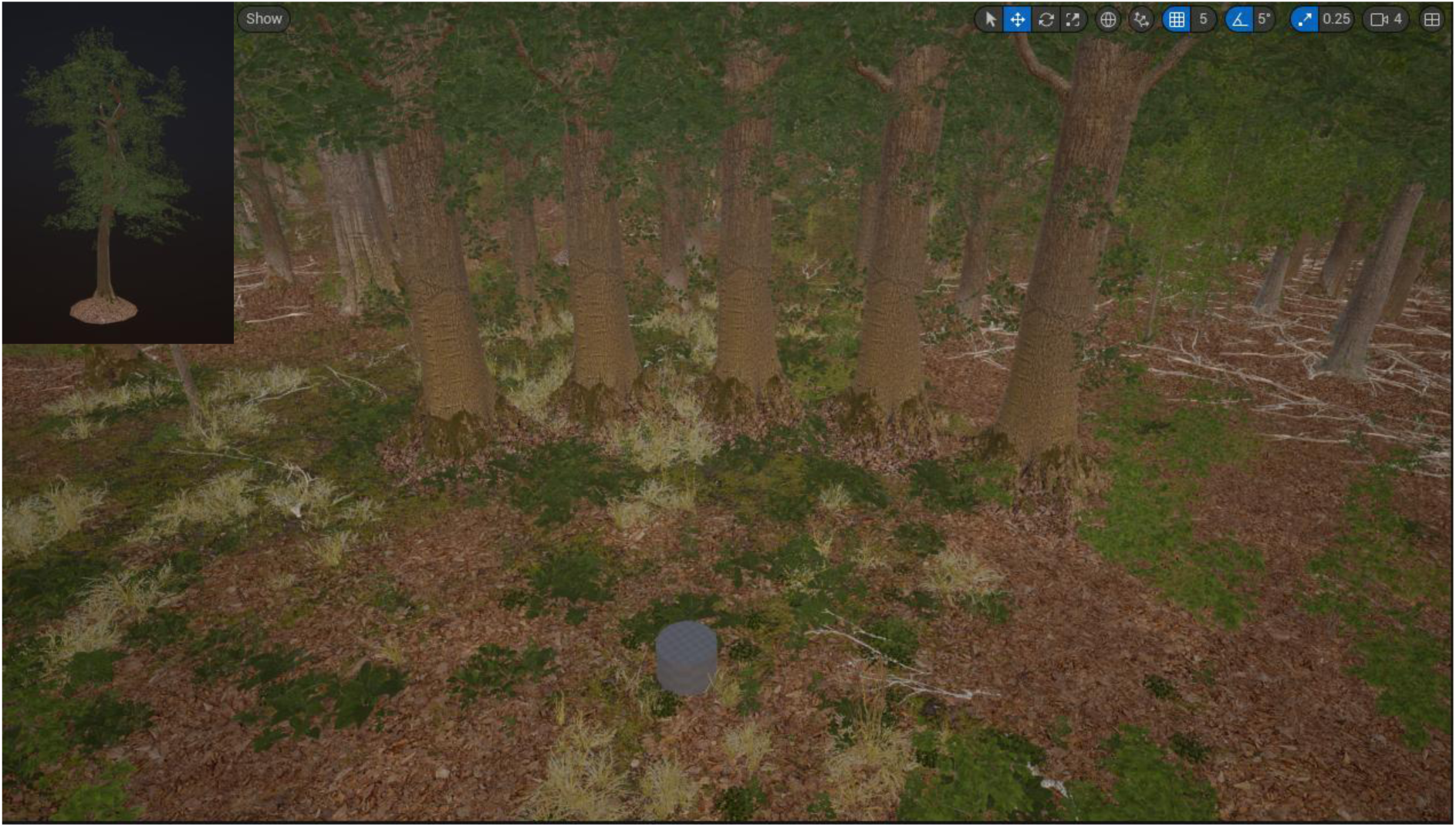
The layout of the tethered environment. Top left: the asset from which the local background is created. Main image: an example of the five local background trees, upon which the target would appear, with the global background behind it. The grey cylinder represents the location of the participant but is not visible during the experiment.

**Supplementary Figure 7.**
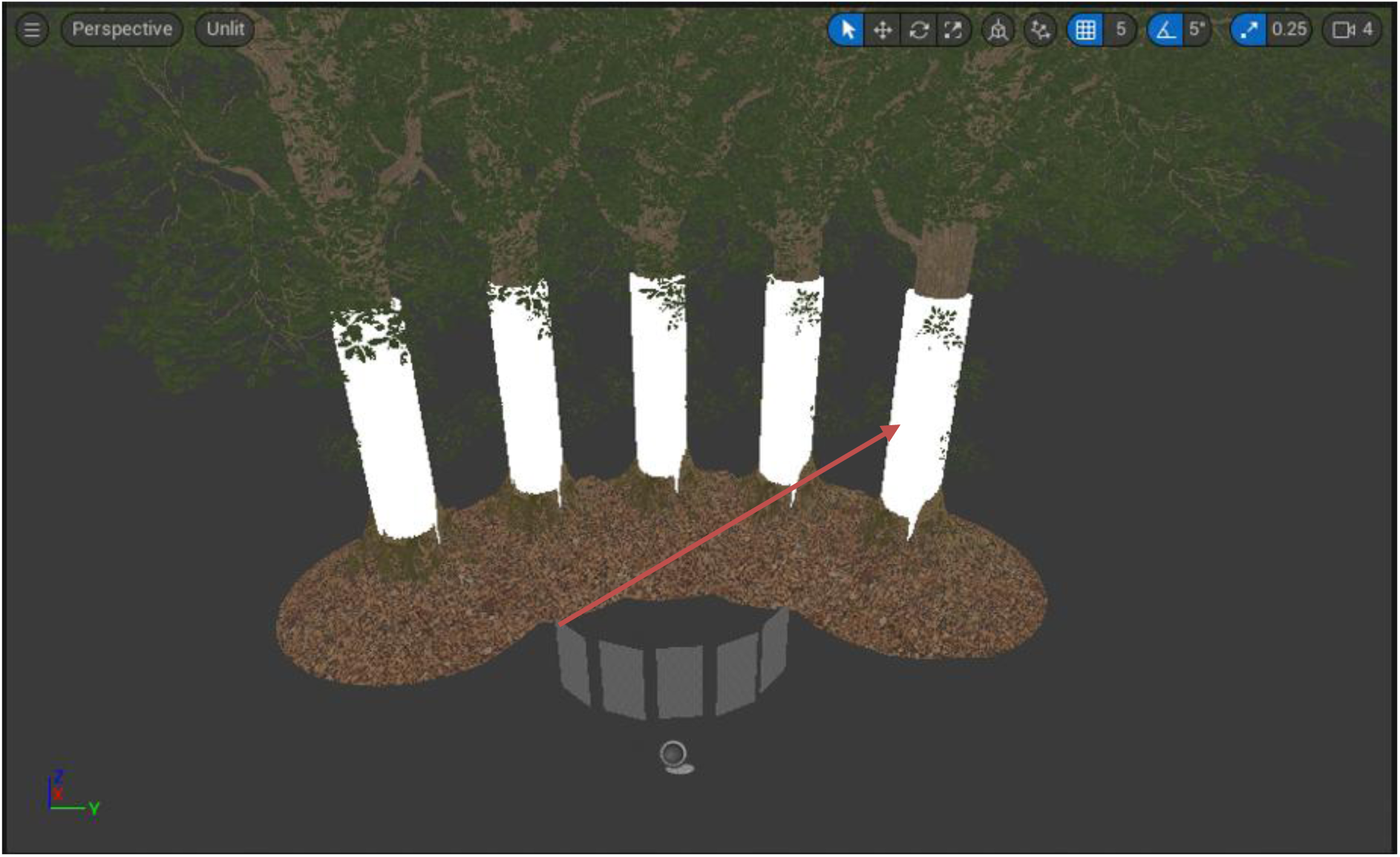
Determining the moth’s location on the local background. Random coordinates are taken in the cuboidal plane (grey) and a line trace (orange) from that coordinate is ‘drawn’ toward the tree. The coordinates at which the line trace contacts the cylinder that encapsulates the tree (white) are the coordinates where the target will appear. This process ensures that each target location on each trial was randomly selected.

**Supplementary Table 1.** The number of predation events for each pattern type, and the time taken, with 95% confidence intervals, for 50% of the targets to be ‘predated’.

|  | n | events | median | L95% | U95% |
| --- | --- | --- | --- | --- | --- |
| Pattern = Uniform | 400 | 394 (98.50%) | 4.91 | 4.44 | 5.36 |
| Pattern = DCHi | 400 | 266 (66.50%) | 12.48 | 11.06 | 15.71 |
| Pattern = DCHiEEHi | 400 | 305 (76.25%) | 8.91 | 7.81 | 10.28 |
| Pattern = DCHiEELo | 400 | 278 (69.50%) | 12.59 | 10.56 | 14.14 |
| Pattern = DCLo | 400 | 275 (68.75%) | 9.92 | 8.66 | 13.64 |
| Pattern = DCLoEEHi | 400 | 198 (49.50%) | NA | 25.83 | NA |
| Pattern = DCLoEELo | 400 | 220 (55.00%) | 23.24 | 16.98 | NA |
| Pattern = EEHi | 400 | 350 (87.50%) | 6.27 | 5.42 | 7.08 |
| Pattern = EELo | 400 | 386 (96.50%) | 5.05 | 4.64 | 5.49 |

**Supplementary Table 2.**
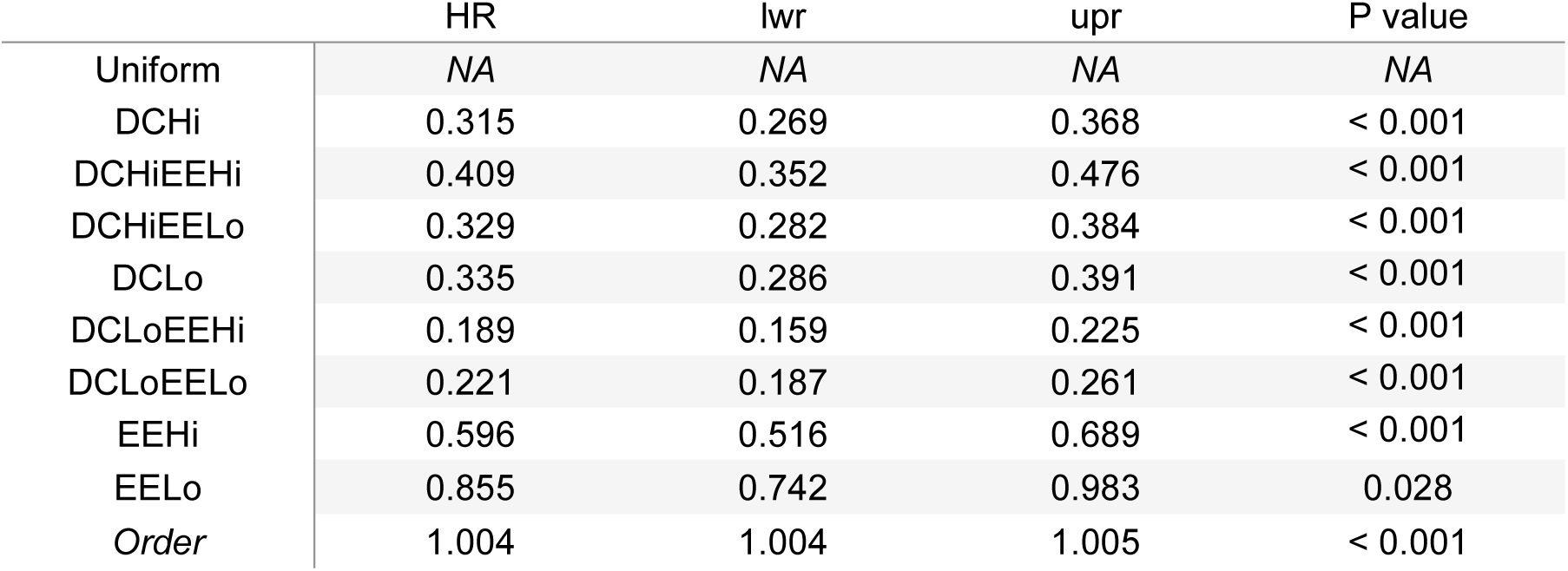
The hazard ratios (HRs) of the fixed effects in the refitted mixed-effects cox model. Note that the hazard ratios of each pattern condition are in relation to Uniform patterning.

**Supplementary Table 3.** The odds ratios (ORs) of the pattern conditions in relation to Uniform patterning.

| Pattern | OR | lwr | upr | $\chi^2$ |
| --- | --- | --- | --- | --- |
| **Uniform** | 1.000 | NA | NA | NA |
| **DCHi** | 32.166 | 15.217 | 83.669 | < 0.001 |
| **DCHiEEHi** | 19.907 | 9.342 | 52.119 | < 0.001 |
| **DCHiEELo** | 28.029 | 13.236 | 72.999 | < 0.001 |
| **DCLo** | 29.029 | 13.716 | 75.636 | < 0.001 |
| **DCLoEEHi** | 65.046 | 30.895 | 168.117 | < 0.001 |
| **DCLoEELo** | 52.192 | 24.780 | 135.526 | < 0.001 |
| **EEHi** | 9.149 | 4.176 | 24.336 | < 0.001 |
| **EELo** | 2.344 | 0.921 | 6.793 | 0.070 |

